# Local Oxygen Tension Dictates Hematopoietic Cell Growth and Potency

**DOI:** 10.1101/2025.09.18.674068

**Authors:** James Ropa, Sarah Gutch, Lindsay Wathen, Gracie Whitacre, Arafat Aljoufi, Scott Cooper, Wouter Van’t Hof, Mark H. Kaplan, Maegan L. Capitano

## Abstract

Hematopoietic stem and progenitor cells support a lifetime supply of blood and immune cells, can become neoplastic when dysregulated, and constitute a powerful cell therapy vehicle for hematologic diseases. Here we provide the most comprehensive study of hematopoietic oxygen (O_2_) dependency to date, demonstrating that human hematopoietic cell numbers, growth, biochemical properties, and functional potency is affected by variation in physiologically and clinically relevant local O_2_ tensions. Lineage defined progenitor cells showed increased expansion in high oxygen, while primitive cells and those with *in vivo* potency were maintained at higher frequencies in low physiologic O_2_. We also present a novel hematopoietic cell oxygen-dependent single cell transcriptomic profile. This and biochemical validation revealed that low O_2_ preserves cells with lower metabolic activity in a less proliferative state that exhibit decreased accumulation of stress markers. Transcriptomics and mouse modeling also elucidated oxygen-dependent mRNA markers of hematopoietic potency. These data reveal oxygen-sensing pathways as targets to improve hematopoietic cell therapies and suggest that local O_2_ tension dictates hematopoietic potential in anatomic niches.

## Introduction

Hematopoietic stem (HSCs) and progenitor cells (HPCs) are responsible for consistent replenishment of blood and immune cells throughout human life, are a source of life-saving treatment in the form of cell therapies, and are dysregulated in various diseases including malignancy^1,2^. A comprehensive understanding of intrinsic and extrinsic factors that affect the function and potency of HSCs/HPCs is still desired. Oxygen (O_2_) is an extrinsic environmental factor with the potential to affect cellular function^3^. Different hematopoietic niches exist at variable local O_2_ tensions, which are also hypoxic compared to ambient air^4–9^. Many tissues such as the bone marrow (BM) exhibit gradients of oxygenation within the same niche^4–9^. Understanding how the various O_2_ tensions that HSCs/HPCs are exposed to throughout their lifespan affects their function will provide insights into their regulation and represents a potential tool to modify their function. Acute exposure to very low O_2_ availability (hypoxia) is well known to affect the health of cells^10^. Regional hypoxia has become an area of great interest in the field of tumor biology, as variable O_2_ levels affect tumor growth, metastasis, and responses to therapies^11,12^. Further, work by our group has shown that exposure to extra physiologic O_2_ that occurs when cells are manipulated *ex vivo* negatively impacts HSC function^13–15^. If and how the entire range of physiologic and non-physiologic O_2_ tensions that hematopoietic cells are exposed to affects their growth and function requires further elucidation.

Factors affecting cell potency also have important implications for hematopoietic cell therapies, which utilize hematopoietic cells isolated from BM, mobilized peripheral blood (mPB), or umbilical cord blood (CB) to treat patients with hematologic disorders^16^. For CB in particular, cell potency (the ability of the cells to repopulate a healthy hematopoietic system), is a critical limiting factor for efficacious treatment^17–20^. Improving the potency of HSCs/HPCs with treatments or *ex vivo* expansion may enhance the efficacy of CB transplantation and other hematopoietic cell therapies like gene editing that require *ex vivo* manipulation^21,22^. Expansion of HSCs/HPCs under hypoxia has been studied as a possible way to improve cell potency, but identifying the optimal O_2_ tension and elucidating the heterogeneous effect that varying O_2_ levels has on HSCs/HPCs has remained challenging^23–26^.

In this study, we hypothesized that cells utilize variable local O_2_ tensions to perform distinct functions and examined how a wide range of oxygenation affects the phenotypic, functional, expansion, and molecular characteristics of primary human CD34+ cells, which are enriched for HSCs/HPCs. Our findings can be used to improve hematopoietic cell therapies and provide insight into O_2_ as a critical regulator of hematopoietic cell function in different anatomic niches.

## Methods

### Umbilical Cord Blood Primary Samples

Cord blood units were collected and distributed under the supervision of Cleveland Cord Blood Center. CD34+ cells were enriched within 55 hours of birth using density gradient centrifugation and Miltenyi enrichment kits.

### ex vivo Expansion in Variable Oxygen

5×10^4^ CD34+ cells were plated in SFEM II serum free media supplemented with 100ng/mL human TPO, FLT3L, and SCF. Cultures were incubated at 37°C, 5% CO_2_ and 1%, 3%, 5%, 14%, or 21% O_2_.

### Mouse Model of Transplantation

24 hours prior to transplantation, female NOD-scid IL2Rgamma^null^ (NSG) were sublethally irradiated (350gy). Mice were transplanted by tail vein injection with 2.5×10^4^ freshly thawed unmanipulated CD34+ enriched cells or 2.5×10^4^ CD34+ cells plus all expanded progeny.

### RNA-sequencing

Bulk RNA was harvested using the Qiagen RNeasy Micro Plus Kit. Libraries were prepared using the SMART-Seq v4 Ultra Low Input RNA Kit (Takara) or the NEBNext Ultra Low Input RNA Sequencing Kit (NEB). Single cells expanded in low O_2_ (1%, 3%, and 5%) were harvested in a hypoxia chamber equilibrated to 3% O_2_. Unmanipulated cells and those expanded at high O_2_ tensions (14% and 21%) were harvested ambient air. Cells were hashed by condition. Using the PIPseq T20 kit (Fluent Biosciences), 5×10^4^ cells from each CBU was subjected to single cell barcoding using particle emulsions and libraries were prepared according to manufacturer’s recommendations^27^. Sequencing was performed by the IU Center for Medical Genomics.

### Data and Code Availability

RNA-sequencing results will be deposited in the NCBI Gene Expression Omnibus (GEO) database. Detailed scRNA-seq analyses are available via Mendeley, DOI: 10.17632/8zzm2nr4w6.1.

## Results

### Hematopoietic cells from differently oxygenated niches exhibit distinct properties

Each hematopoietic niche exists at different O_2_ tensions (Table 1)^4–9^. The BM ranges from <1%-6% O_2_ while peripheral blood (PB) total oxygenation lies between 4-14% (with location dependent levels of hemoglobin-bound or dissolved O_2_^28^) and is different in arterial or venous blood. CB exhibits ranges from 0.5-6% O_2_^7,8^, though possibly up to 10% (this study, Figure 1D). Other tissues where HSCs/HPCs reside like the lung are highly oxygenated, up to 17%^9,29^. We infer that HSC/HPC responses to O_2_ could be a consequence of specialization of the tissue they reside in. To test differences in HSCs/HPCs from differently oxygenated niches, we performed *ex vivo* characterization of HSCs/HPCs from human CB, mPB, or BM CD34+ cells using immunophenotypic analysis by cell surface staining with flow cytometry and colony forming unit (CFU) assays (Figure 1A-C)^30,31^. This showed that the HSC/primitive HPC enriched CD34+CD38-fraction is higher in mPB and CB compared to BM (Figure 1A). There was also a significantly higher frequency of multipotent progenitors (MPPs) and multilymphoid progenitors (MLPs) in CB compared to BM (Figure 1A-B). Conversely, BM contains a higher proportion of more lineage determined CD34+CD38+ HPCs, with a significantly higher granulocyte-macrophage progenitors (GMPs) frequency compared to mPB (Figure 1A-B). CB and mPB show increased trends of functional HPCs, demonstrated by numbers of CFU-granulocyte, macrophage (CFU-GM) and CFU-granulocyte, erythroid, macrophage, megakaryocyte (CFU-GEMM) compared to BM (Figure 1B-C). These data suggest that CD34+ cells exhibit different proportions of HSCs/HPC subpopulations when isolated from different niches.

**Figure 1.**
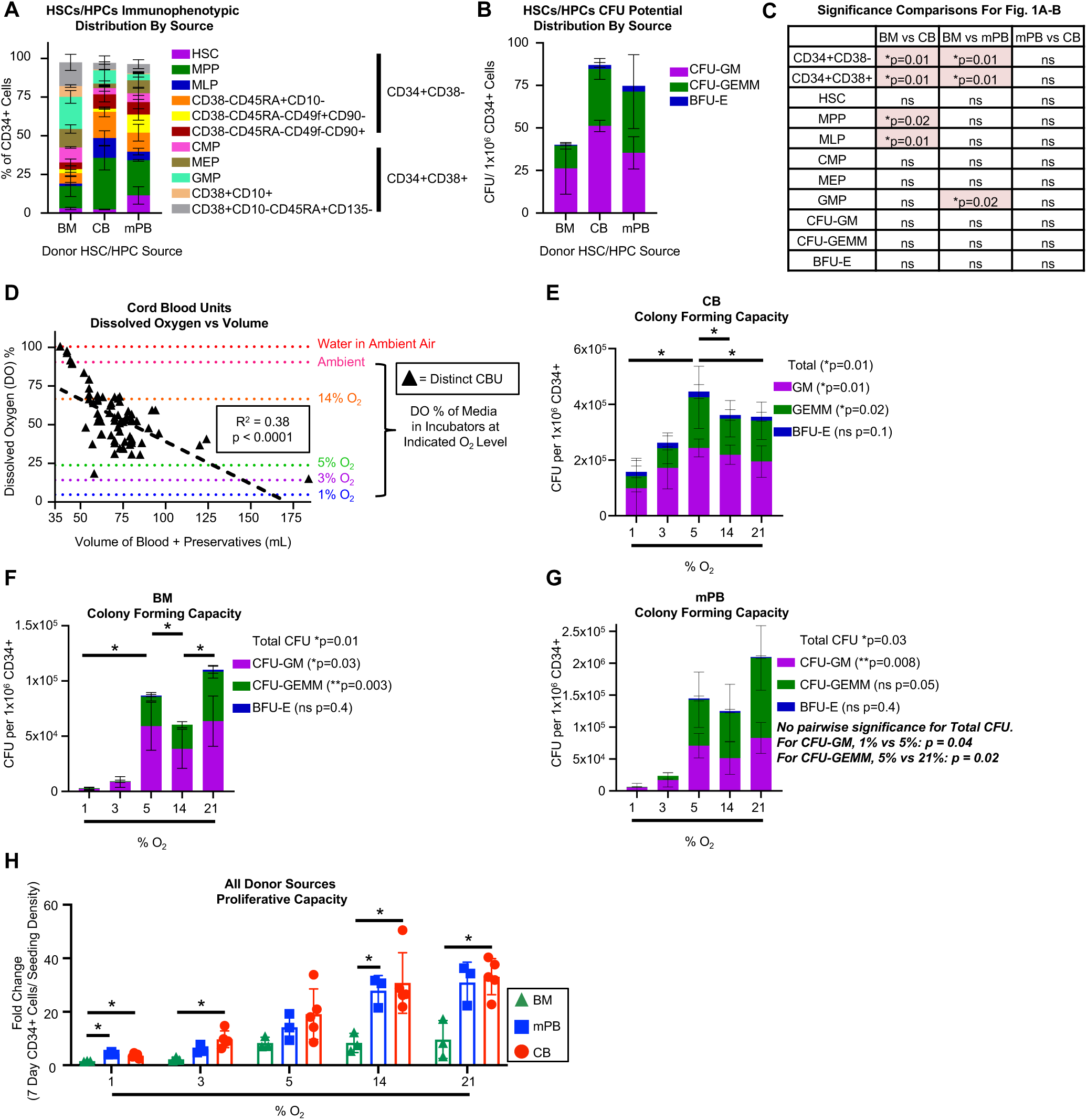
HSCs/HPCs display O_2_ dependencies. A) Immunophenotypically defined HSCs/HPCs or B) colony forming unit (CFU) numbers as a fraction of CD34+ cells from bone marrow (BM, n=3), mobilized peripheral blood (mPB, n=3), and cord blood (CB, n=2). C) Significance analysis of A using 1-way ANOVA with post hoc comparisons. D) Dissolved O_2_ concentration in fresh whole CBUs compared to total volume of the CBU. Dissolved O_2_ levels associated with physiologically relevant local and atmospheric O_2_ tensions are indicated by dashed lines. E-G) CFU from CB (n=8), BM (n=3), or mPB (n=3) CD34+ cells performed in the indicated O_2_ tensions. 12 days after plating CFUs of different potentials were enumerated manually by counting. H) CB (n=5), BM (n=3), or mPB (n=3) CD34+ cells expanded in serum free media with growth factors for 7 days in the indicated O_2_ tensions. Total CD34+ cell numbers compared to the number of cells seeded by flow cytometry analysis. Stats: one-way ANOVA controlling for donor unit (indicated by p-value) with post hoc Tukey or Sidak comparisons. *p<0.05.

**Table 1.**
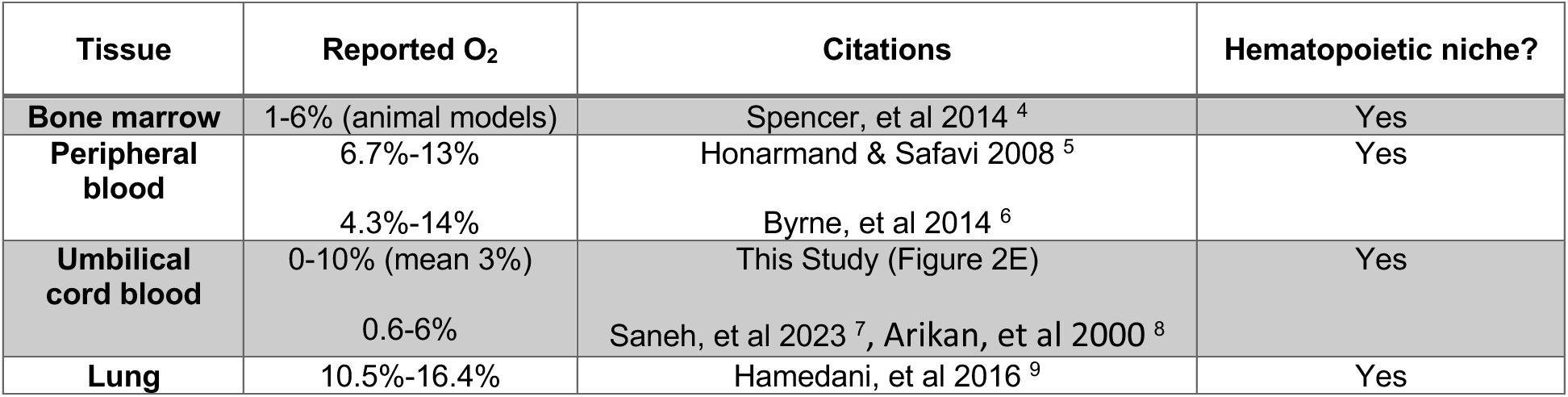
O_2_ Tensions of Hematopoietic Tissues. Shown are modelled or measured O_2_ tensions in various hematopoietic niches as reported in the literature. The values from “This Study” were calculated based on the dissolved O_2_ content of umbilical cord blood, accounting for dilution of the cord blood hypoxic state that occurs when preservatives are added to the total blood volume (Figure 1D).

We have previously suggested that donor hematopoietic cells used in therapy and research are necessarily exposed to ambient O_2_ levels upon harvest and thus lose their relative hypoxic states^13–15,32^. To test this directly, we measured the dissolved O_2_ concentration in > 70 fresh CBUs that were never frozen and were measured within 1 week of isolation. To our surprise, the dissolved O_2_ level of CBUs collected in preservation bags remains hypoxic relative to fully oxygenated liquids (Figure 1D). The level of dissolved O_2_ directly correlates with the volume of the unit. This is likely the result of the physiologic oxygenation state of the blood being slightly diluted by the addition of fully oxygenated anti-coagulant preservatives in each CBU, which impacts low collection volumes the most. Thus, CB remains lowly oxygenated after collection. We thus focus on CB as a primary model for studying O_2_ dependency. We also examine BM and mPB to explore differential O_2_ sensitivity of HSCs/HPCs from different donor sources, inferring that their exposure to extra physiologic O_2_ is likely similar to that of CB.

### Hematopoietic cells exhibit O_2_ dependency *ex vivo*

Differently oxygenated hematopoietic niches have non-O_2_ related environmental factors that could account for changes to cellular properties. We thus sought to test the effects of O_2_ on hematopoietic cell growth and function in a single variable system using oxygen-controlled incubators. We maintained CD34+ cells in 1%, 3%, 5%, 14%, and 21% O_2_, which spans the range of O_2_ levels in tissues in which hematopoietic cells reside, migrate through, or are processed (Table 1). First, we measured the CFU potential of the CD34+ cell fraction and found that total CFU, CFU-GM, and CFU-GEMM numbers from CB, BM, and mPB are significantly altered by local O_2_ tension. Total CB CFU numbers are significantly higher in 5% O_2_ compared to 14% and 21% and CB CFU-GM numbers are higher in 5% compared to 1% (Figure 1E), while BM and mPB CFU numbers show direct correlation to O_2_ tension, with increased CFU-GM, CFU-GEMM, and total CFU at 5%, 14%, and 21% O_2_ compared to lower tensions (Figure 1F-G). Burst forming unit-erythroid (BFU-E) numbers were unaffected by variable O_2_. Next, we cultured CD34+ cells in serum free media with growth factors for 7 days to measure their proliferative capacity. CB and mPB CD34+ cells exhibit increased proliferation with increased O_2_ tension (Figure 1H). Interestingly, at all tensions except 5% O_2_, CD34+ cells from mPB and/or CB exhibited significantly higher proliferation compared to BM.

### Multipotent hematopoietic cells exhibit preferential expansion in different O_2_ ranges compared to lineage determined HPCs

CD34+ cells are heterogenous, so we next used immunophenotypic analysis following culture to determine how different HSC/HPC subpopulations expand in variable O_2_. For cell fractions derived from CB and mPB but not BM, total nucleated cells, total CD34+ cells and the multipotent cell enriched CD34+CD38-fraction, and the lineage defined progenitor enriched CD34+CD38+ fraction exhibited significantly increased expansion in increased O_2_ with higher total cell numbers at 5%, 14%, and 21% O_2_ compared to 1%, 3% and input (Figure 2A-C; Figure S1A-I). HSC growth was significantly affected by O_2_ tension with peak expanded cell numbers in 5% O_2_ in cells from mPB and BM, while HSCs from CB exhibited a similar trend (Figure 1D-F). MPPs from all sources are also sensitive to changing O_2_, but the ideal growth oxygenation varies between CB, BM, and mPB (Figure 2G-I). Interestingly, when combining all donor sources HSCs exhibit the highest frequency of HSCs in 5% O_2_, while MPP frequencies are highest in 1-3%, suggesting these cell types preferentially proliferate in different local O_2_ levels (Figure 2J-K)

**Figure 2.**
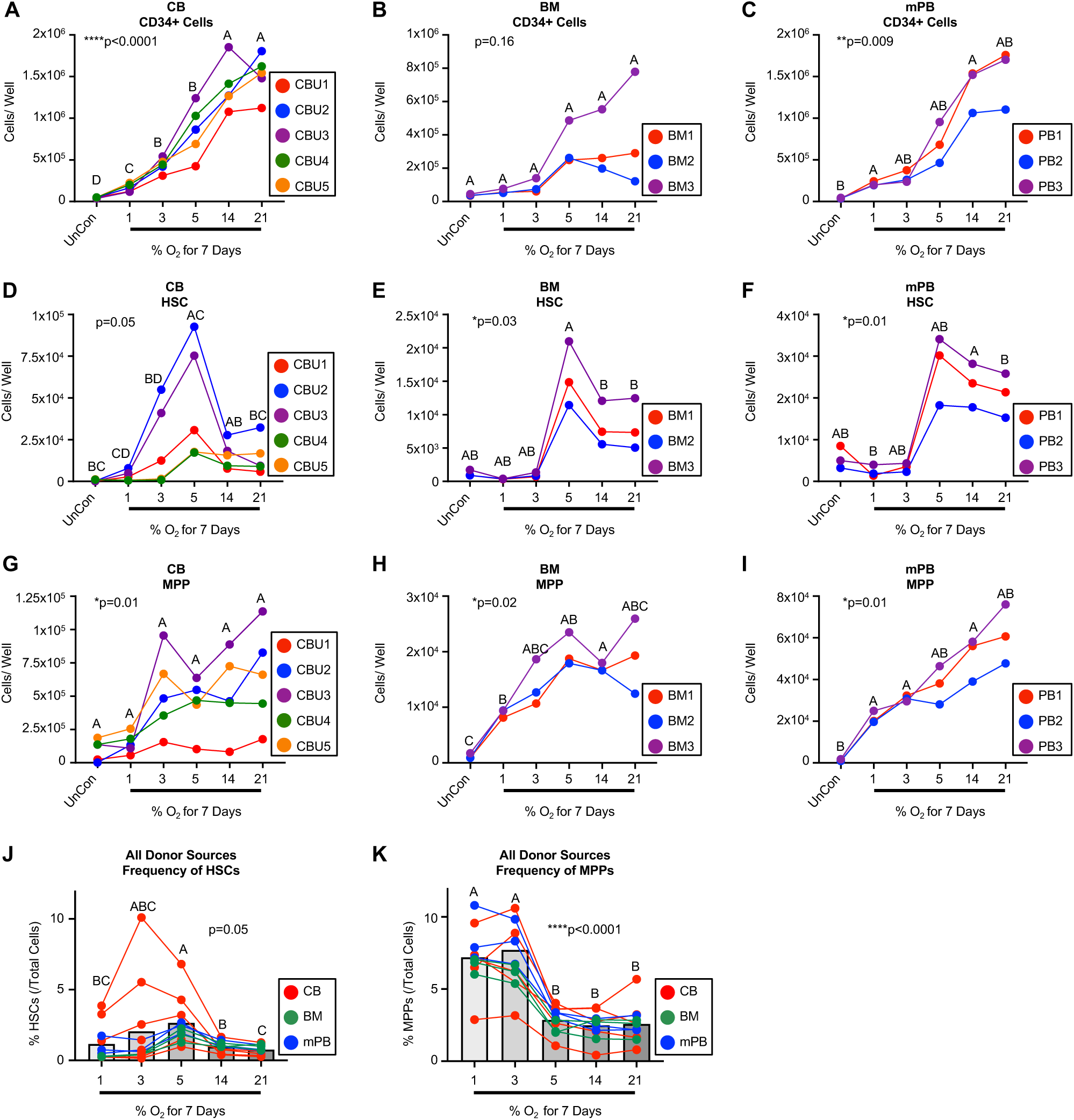
Multipotent HSCs/MPPs expand the most in low physiologic O_2_ tension. A-I) Cord blood (CB n=5), bone marrow (BM n=3), or mobilized peripheral blood (mPB n=3) CD34+ cells were expanded in serum free media with growth factors for 7 days in the indicated oxygen tensions. Cells were then analyzed for enumeration of total numbers of the indicated immunophenotypically defined population by flow cytometry. J-K) Percent of indicated immunophenotypic HSC/HPC population as a fraction of total cells in the well after 7 days of expansion. Stats: one-way ANOVA (indicated by p-value) with post hoc comparisons indicated by compact letter display (in compact letter display, groups that are not statistically different are indicated by matching letters). Each point/color indicates distinct biological donor sources as indicated by the included legends. UnCon = Unmanipulated control (input).

Lineage defined progenitors including MLPs, GMPs, and CMPs expanded significantly more in high O_2_ tensions with CB showing the significantly higher expansion in 5-21% compared to lower tensions with similar trends in BM and mPB (Figure 3A-F, Figure S1J-L). Megakaryocyte-erythroid progenitor (MEP) expansion was not significantly affected by O_2_ tension (Figure S1M-O). In combining all donor sources, frequencies of MLPs were lowest in 5% O_2_ but similar in all other tensions, while GMP proportions were highest in 14-21% O_2_, showing still different O_2_ dependencies from HSCs/MPPs (Figure 3G-H). To test the function of these expanded cells, we performed CFU assays on CD34+ cells expanded from CB. CFU-GM and CFU-GEMM both expanded more in liquid culture at 5%, 14%, and 21% compared to lower O_2_ tensions (Figure 3I-J). Together, these data show that immunophenotypic mature progenitors and functional CFUs exhibit increased expansion in higher O_2_ levels, though responses vary slightly with donor source. Importantly, HSCs and more primitive HPCs maintain higher proportional representation in the expanded pool of cells in low physiologic O_2_, though this varies by cell type.

**Figure 3.**
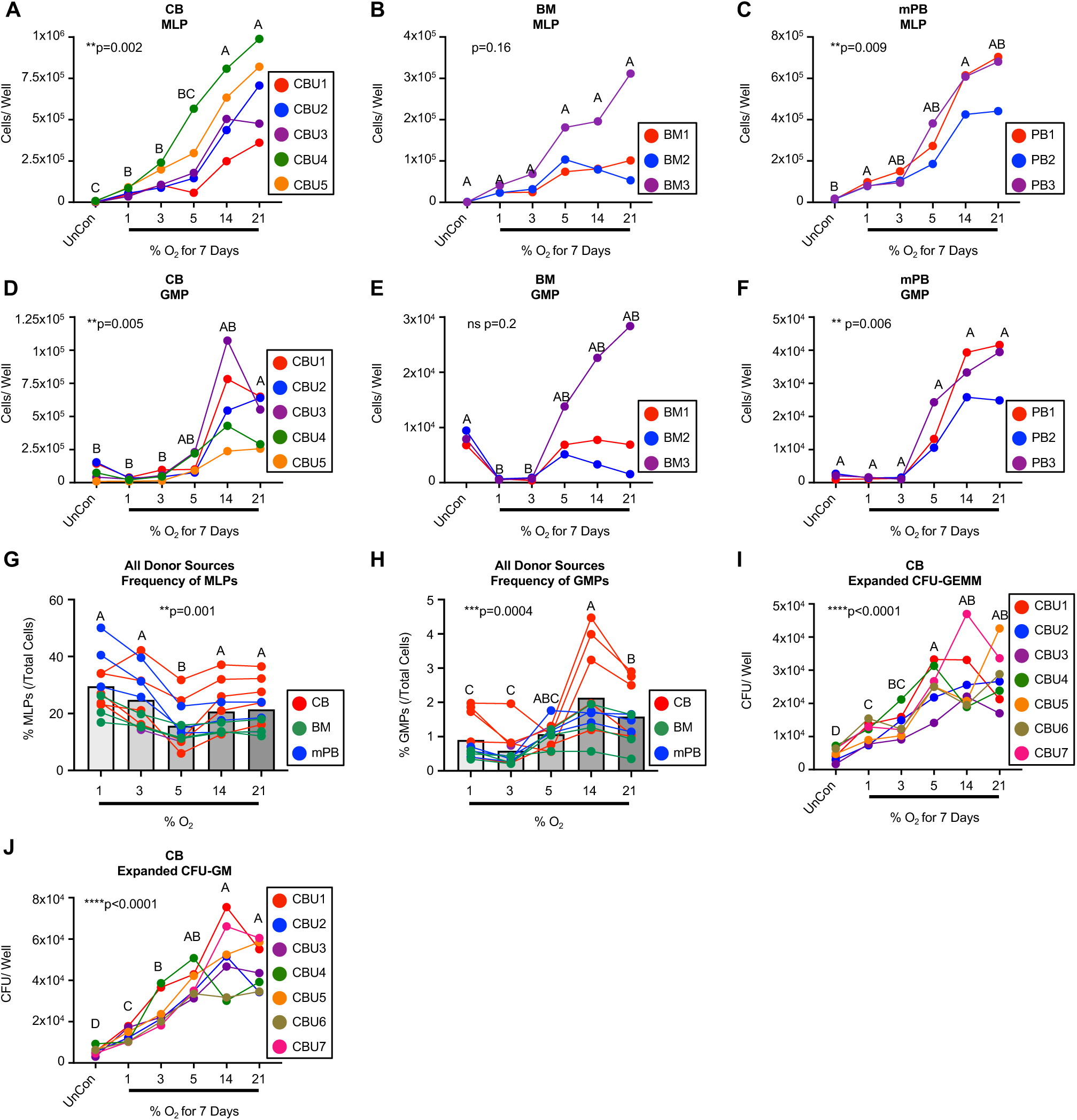
Lineage committed HPCs exhibit differential preferences for O_2_ tension. A-F) Cord blood (CB n=5), bone marrow (BM n=3), or mobilized peripheral blood (mPB n=3) CD34+ cells were expanded in serum free media with growth factors for 7 days in the indicated oxygen tensions. Cells were then analyzed for enumeration of total numbers of the indicated immunophenotypically defined population by flow cytometry. G-H) Percent of indicated immunophenotypic HPC population as a fraction of total cells in the well after 7 days of expansion. I-J) CB CD34+ cells (n=7) were expanded in serum free media with growth factors for 7 days in the indicated oxygen tensions. Cells were then plated in CFU assays at the same tension (5%). 12 days after plating CFUs of different potentials were enumerated manually by counting. Stats: one-way ANOVA (indicated by p-value) with post hoc comparisons indicated by compact letter display (in compact letter display, groups that are not statistically different are indicated by matching letters). Each point/color indicates distinct biological donor sources as indicated by the included legends. UnCon = Unmanipulated control (input).

### CD34+ expansion in variable O_2_ affects *in vivo* hematopoietic engraftment

To determine whether local O_2_ tensions affect the *in vivo* potency and engraftment kinetics of HSCs/HPCs, we examined the ability of cells expanded at variable O_2_ levels to repopulate a hematopoietic system *in vivo* (Figure 4A). CD34+ cells were expanded in five O_2_ tensions for 7 days in serum free media with growth factors. Unmanipulated “input” cells (2.5×10^4^ cells) or input cells plus their expanded progeny (variable numbers based on total cell expansion) were transplanted to NSG mice. While total human engraftment was not significantly different at 2 weeks post transplantation (Figure 4B), early human neutrophil recovery measured in mouse PB, an analogous measure of successful patient transplantation, was increased when cells were expanded in low physiologic O_2_ tensions (1-5%) compared to input cells (Figure 4C). When correcting for differential cell numbers transplanted due to differential expansion, early neutrophil recovery as a fraction of transplanted CD34+ cells was significantly higher in 1% O_2_ than in all higher tensions (Figure 4D). 6 weeks post transplantation, mice transplanted with cells expanded at 1% O_2_ showed lower total engraftment in the PB compared to higher tensions (Figure 4E), a transient effect no longer observed by 10 weeks post-transplantation (Figure 4F), but which remained a trend in the BM at 16 weeks post transplantation (Figure 4G). At 10 weeks, mice transplanted with expanded cells at all O_2_ tensions had significantly higher human myeloid/lymphoid ratio in their PB compared to those transplanted with unmanipulated input cells, with mice transplanted with cells expanded at 1% trending toward the most “balanced” ratio (4H). No differences in myeloid/lymphoid balance were observed in the BM 16 weeks post transplantation, indicating either a transient effect or one that can only be observed in the PB (Figure 4I). In secondary engraftment assays, CD34+ cells initially expanded at 14% O_2_ yielded the highest secondary engraftment and was the only condition where all mice had observable secondary engraftment (4J, data not shown). Together, these data show that different cell populations are responsible for different engraftment kinetics, such as early myeloid recovery and long-term reconstitution, and that these subpopulations are variably affected by physiologic levels of O_2_.

**Figure 4.**
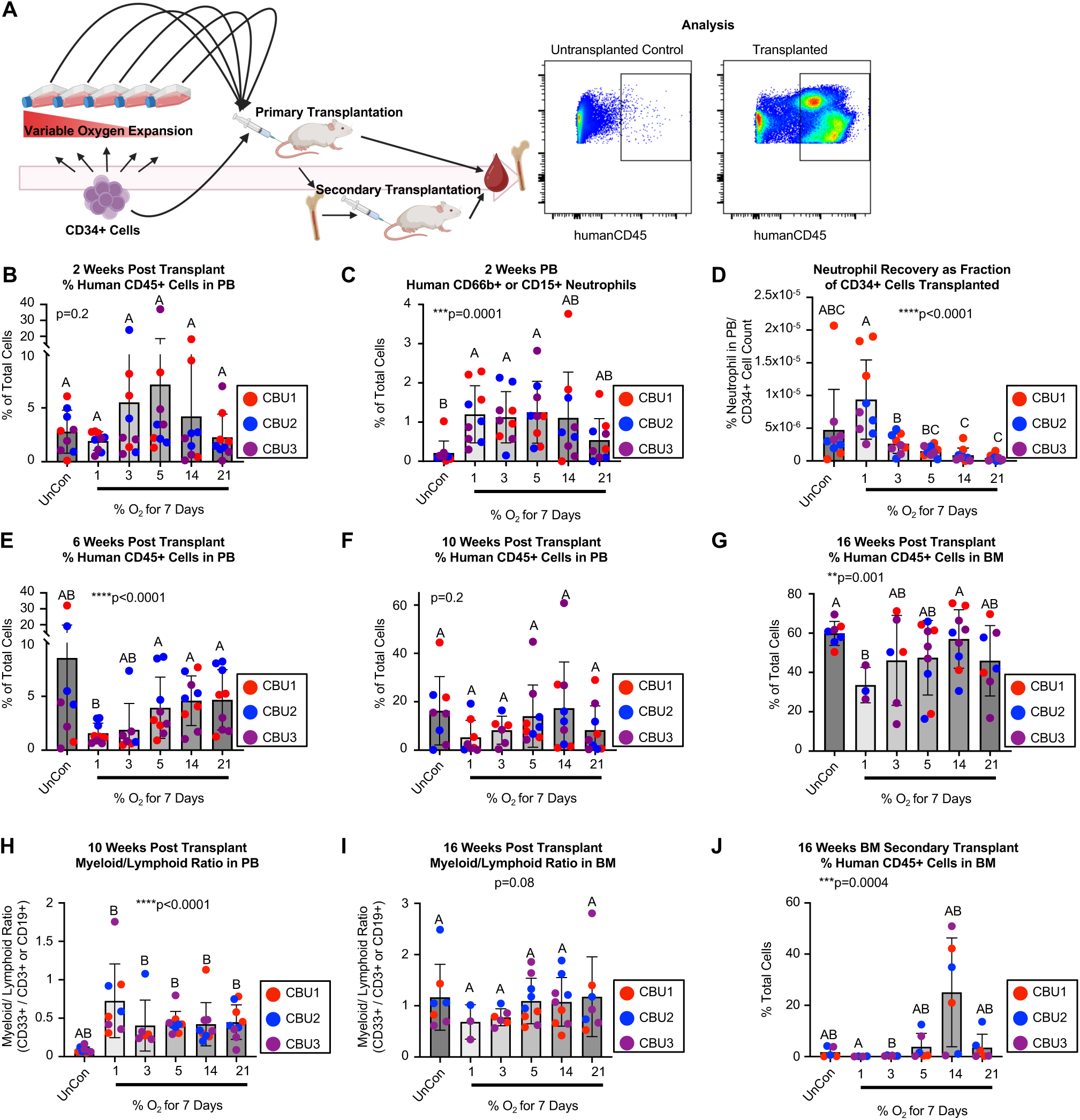
Expanded HSC/HPC *in vivo* potency is affected by local O_2_ tension. A) Experimental scheme for *in vivo* studies. B-J) CB CD34+ (n=3) cells expanded in serum free media with growth factors for 7 days in the indicated O_2_ tensions. Input cells (25,000/ mouse) or input cells plus expanded progeny were transplanted to sublethally irradiated NSG mice (n=9 each group). B-C) Mice were monitored at 2 weeks for total human CD45+ chimerism and neutrophil recovery in PB by flow cytometry. D) Neutrophil recovery percentage as a fraction of total cells transplanted (25,000 input cells + expanded progeny). E-F) Mice were monitored at 6 and 10 weeks for total human CD45+ chimerism in PB by flow cytometry. G) Mice were monitored at 16 weeks for total human CD45+ chimerism in BM by flow cytometry. H-I) Mice were monitored at 10 weeks in the PB and 16 weeks in the BM for myeloid/lymphoid recovery by flow cytometry. Myeloid cells were defined by CD33+ cells while lymphoid cells were positive for CD3 and/or CD19. J) Pooled BM cells from the primary transplant recipients were injected into sublethally irradiated secondary recipients (n=5 each group). Engraftment was monitored at 16 weeks post transplantation in the BM of secondary recipients. Stats: one-way ANOVA (indicated by p-value) with post hoc comparisons indicated by compact letter display (in compact letter display, groups that are not statistically different are indicated by matching letters). Each color indicates distinct biological donor sources as indicated by the included legends. Each point indicates a recipient mouse. Notes on mouse numbers: the following groups lost mice due to experimental conditions (likely bone marrow failure or infection) during the course of the transplantation experiment: 1%-1 mouse; 3%-3 mice; 5%-0 mice; 14%-0 mice; 21%-2 mice. 5 mice were lost from the 1% group due to husbandry complications unrelated to the experimental conditions. UnCon = Unmanipulated control (input).

### Expansion in variable O_2_ alters the hematopoietic transcriptome

We next sought to elucidate mechanisms underlying O_2_ hematopoietic dependency by molecular profiling of CB HSCs/HPCs at a single cell resolution. This would allow us to identify subclusters of cells associated with functional potency and determine the effects of local O_2_ tensions on various subpopulations of HSCs/HPCs. We performed single cell RNA sequencing (scRNA-seq) using particle-templated instant partition sequencing (PIP-seq) to create a novel oxygen-dependent transcriptomic atlas of human hematopoietic cells^27^. Two CBUs were expanded for 7 days at 5 O_2_ levels or were sequenced unmanipulated as input cells. Two additional units were expanded for only 2 days to capture early responses to variable O_2_. Single cell transcriptomes were analyzed using single cell differential expression combined with pseudobulk expression analysis. Cells were clustered and manually annotated in a semi-supervised manner (Figure 5A). As expected, the majority (60%) of unmanipulated CD34+ cells were transcriptomically distinct from cultured cells (Figure 5A, Figure S2A, Table S1) and were separated and sub-clustered for higher resolution (Figure 5A, Figure S2B).

**Figure 5.**
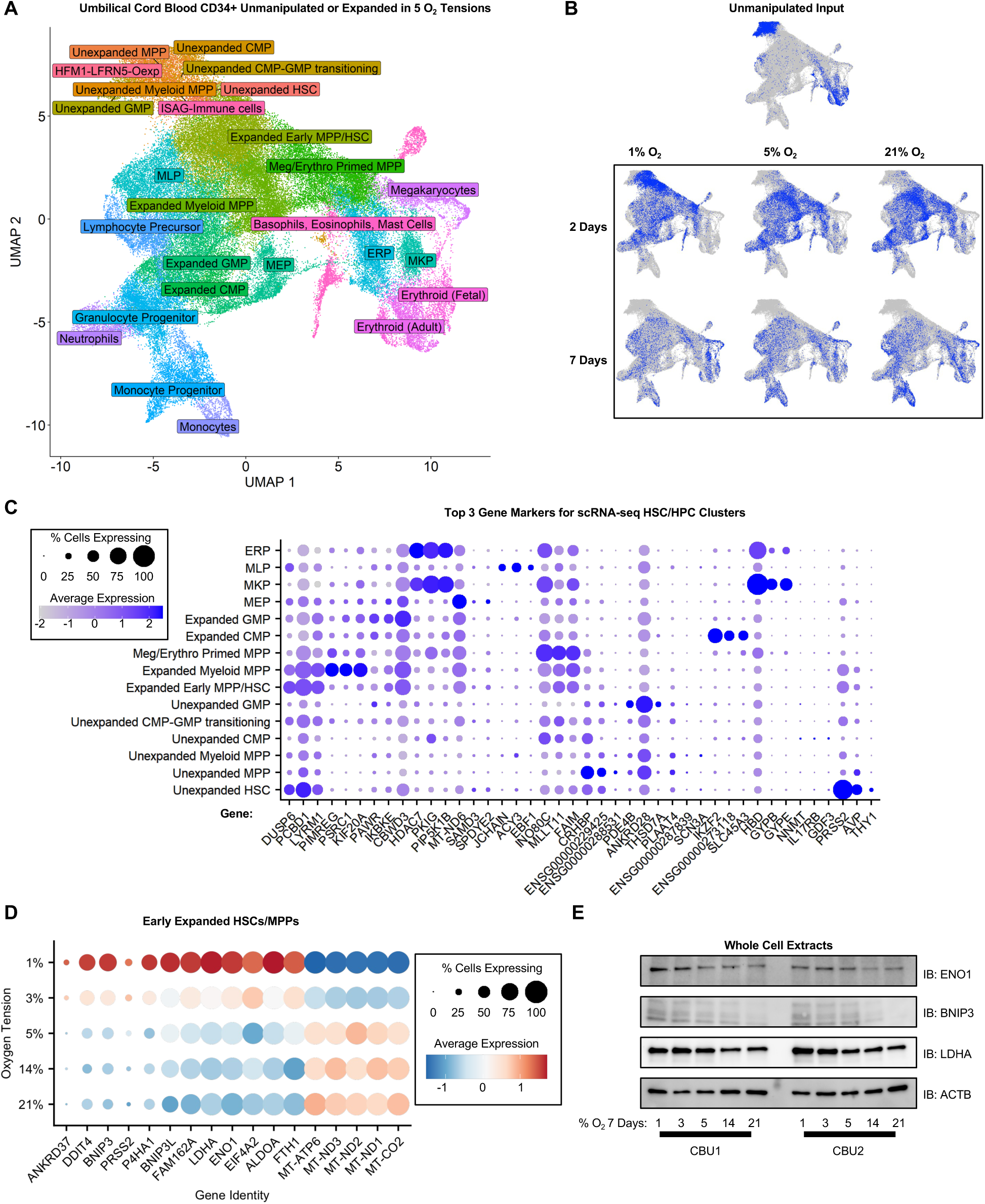
scRNA-seq of HSCs/HPCs reveals markers of potent HSCs/MPPs. A-D) CB CD34+ cells expanded in serum free media with growth factors for 2 days (n=2) or 7 days (n=2) in the indicated O_2_ tensions, or unmanipulated input cells (n=2) were subjected to scRNA-seq using particle-templated instant partition sequencing (PIP-seq). A) UMAP showing annotated clusters from all cells and conditions. B) UMAP plots with cells from the indicated expansion condition highlighted. C) Dot Plot showing top 3 gene markers used to annotate HSC/HPC clusters. D) Dot plot showing genes that are up and downregulated in low and high O_2_ tensions in the Early Expanded HSCs/MPPs subcluster of cells. E) Western blot of whole cell lysates using antibodies targeted against indicated proteins after expansion in the shown O_2_ conditions (IB=immunoblot).

Our final single cell object consists of 26 manually annotated clusters of cells characterized by distinct expression of genes associated with specific hematopoietic cell types (Figure 5A, Figure S3, Table S2). Within our final single cell object we annotated clusters comprised primarily of unmanipulated CBUs and termed them “unexpanded” HSCs, MPPs, myeloid MPPs, CMPs, GMPs, and CMP-GMP transitioning. Other than unmanipulated input, only cells expanded at 1% O_2_ for 2 days maintained “unexpanded” transcriptomic characteristics (Figure 5B). By day 7 of expansion, few cells in any expanded group were found in these clusters, though low O_2_ tensions still maintained higher a higher proportion of cells with these molecular characteristics (Table S2). This confirms that *ex vivo* expansion of CD34+ cells dramatically alters their transcriptomic profile, but this can be mitigated for short time periods by maintaining the cells at very low physiologic (1%) O_2_. Next, we annotated cells enriched for primitive HSC/MPP markers in expanded conditions on day 2 or day 7 and termed them “expanded early HSC/MPP” (Figure 5C). Notably, CD34+ cells expanded at lower O_2_ tensions (1-5%) exhibit a higher proportion of these cells after the full seven days of expansion compared to input and higher O_2_ levels (Table S2).

We inferred based on the known phenomena of HSCs/HPCs losing overall engraftment potential during long-term *ex vivo* culture that “unexpanded” HSC/HPC clusters are enriched with functionally potent cells. We thus analyzed gene markers that define the HSC subpopulation from these cells relative to other cell clusters (Figure 5C; Figure S3). Unexpanded HSCs were characterized by expression of *THY1* and *AVP*, known markers of HSCs in single cell transcriptomics (Figure 5C)^33,34^. Interestingly, these cells also exhibited high expression of *PRSS2*, which is associated with pancreatic functions and does not have a well characterized hematopoietic function^35,36^. *PRSS2* expression is also primarily confined to additional unexpanded cell clusters as well as “Expanded Early HSCs/MPPs” cluster, which we predict contains cells responsible for engraftment from expansion conditions. Our *in vivo* data also leads us to infer that local O_2_ affects HSC/HPC potency, particularly in cells responsible for early immune reconstitution and those that retain a more balanced myeloid/lymphoid ratio. We thus examined genes that are altered based on different local O_2_ tension. We focused this analysis on the “Expanded Early HSCs/MPPs” cluster as a main cell population of interest contains sufficient numbers of cells from each condition for analysis. This primitive HSC/HPC subpopulation showed oxygen-dependent expression of genes associated with regulating metabolism, mitochondrial function, and proliferation. Some of these genes are associated with the mTORc pathway including *BNIP3*, *DDIT4*, *ENO1*, and *LDHA* (Figure 5D)^37–41^. Others are a core group of mitochondrial genes including *MT-ATP6*, *MT-ND1/2/3*, and *MT-CO*_2_ (Figure 5D). *PRSS2*, the marker for unexpanded HSCs, is also shown to be oxygen-dependent in this cell subpopulation (Figure 5D). These genes exhibit O_2_-dose-dependent responses up to 5%, but at 5% O_2_ and above, they show similar expression patterns. We can infer from these data that there is a molecular switch in the low physiologic O_2_ space somewhere between 3%-5% O_2_ responsible for oxygen-dependent gene expression regulation. We confirmed changes in expression of select O_2_ dependent genes including ENO1, BNIP3, and LDHA at the protein level by western blot (Figure 5E). All together, these analyses reveal O_2_ dependent genes associated with potent subpopulations of cells that may be important for HSC/HPC functions.

### O_2_ affects hematopoietic cell stress

We next performed gene set enrichment analyses, comparing the single cell transcriptomic profiles of early expanded HSCs/MPPs grown in 21% compared to 1%. We found that cells expanded in 21% had transcriptomes enriched for genes associated with oxidative stress and mitochondrial function, suggesting an underlying accumulation of acute stress in cells grown at higher O_2_ tensions (Figure 6A-B). Additionally, cells grown at lower O_2_ levels are predicted by their transcriptomes to have a higher proportion of cells in G1 (Figure 6C), which may indicate quiescence.

**Figure 6.**
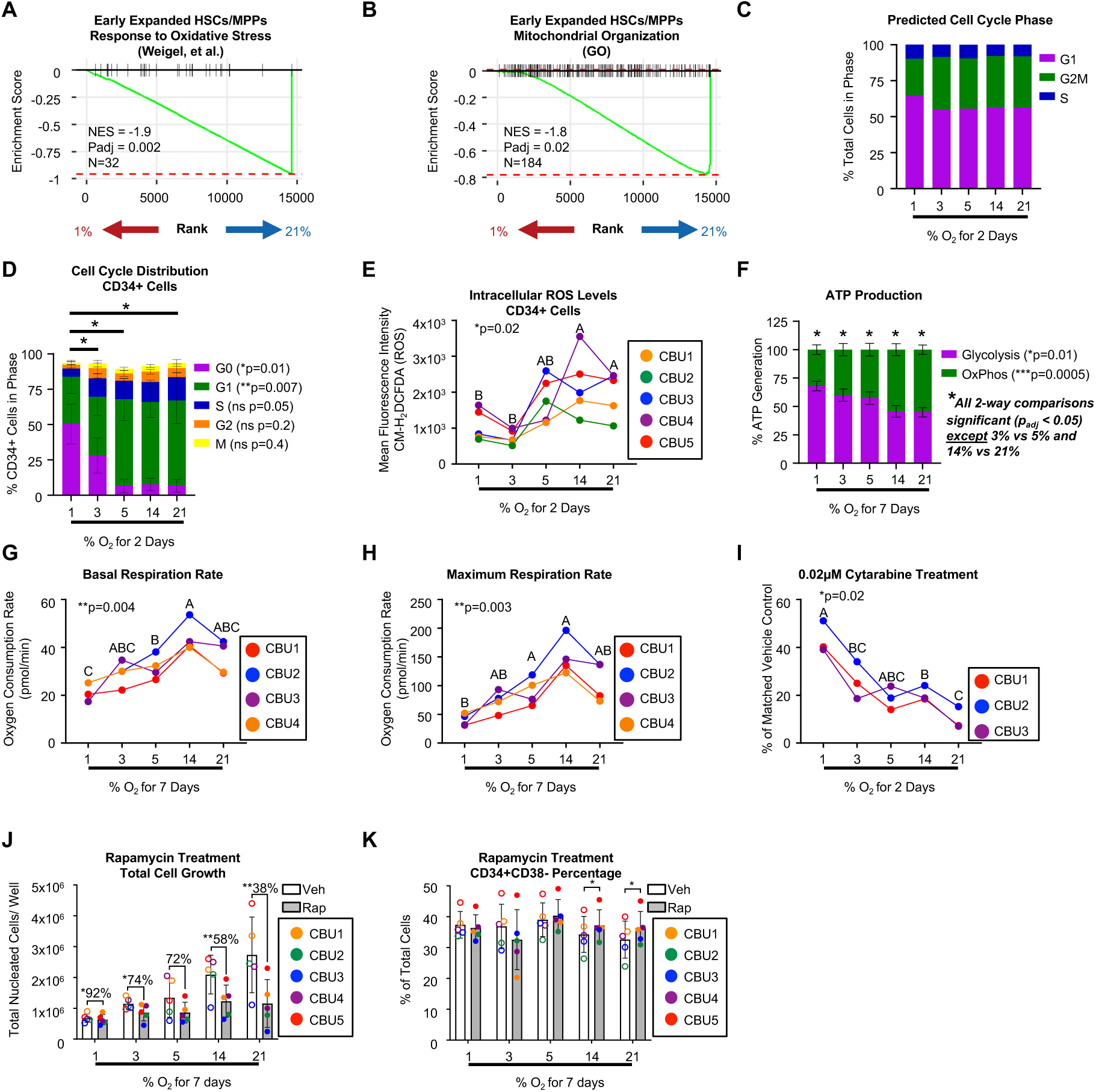
HSCs/HPCs exhibit oxidative stress associated properties in higher O_2_ tensions. A-C) CB CD34+ cells expanded in serum free media with growth factors for 2 days (n=2) or 7 days (n=2) in the indicated O_2_ tensions, or unmanipulated input cells (n=2) were subjected to scRNA-seq using particle-templated instant partition sequencing (PIP-seq). A-B) GSEA plots showing the enrichment for the indicated gene programs in 1% versus 21% O_2_ in the subcluster Early Expanded HSCs/MPPs. C) Transcriptomic prediction of cell cycle status for all cells in the 2-day expanded groups for the indicated O_2_ tensions. D-K) CB CD34+ cells expanded in serum free media with growth factors for 2 or 7 days in the indicated O_2_ tensions. D) Cell cycle analysis using Ki67/DAPI flow cytometry analysis after 2 days expansion (n=3). E) Intracellular reactive O_2_ species (ROS) measurement after 2 days expansion using CM-H2DCFDA ROS indicator and flow cytometry analysis (n=5). F-H) Seahorse metabolic flux analysis after 7 days expansion using F) ATP rate assay to show ATP production (n=4) or G-H) mitochondrial stress test (n=4) to show calculated basal and maximum respiration rates. I) CD34+ cells were treated with 0.02µM cytarabine (n=3) for 2 days in variable O_2_ tensions and counted by Trypan Blue viability staining. J) Total nucleated cells and K) CD34+CD38-cells were enumerated by flow cytometry after 7 days of treatment with 50nM Rapamycin in the indicated O_2_ tensions. Stats: one-way ANOVA (indicated by p-value) with post hoc comparisons indicated by compact letter display (in compact letter display, groups that are not statistically different are indicated by matching letters). Colors used are described in figure legends.

Based on these findings, we tested whether hematopoietic cells exhibit changes in stress associated properties in different local O_2_ tensions. By flow cytometry, we analyzed the cell cycle status of CD34+ cells grown in variable O_2_ tensions. Cells grown in 1% and 3% O_2_ for 2 days had increased proportions in G0 compared to higher O_2_ tensions (Figure 6D), confirming the transcriptomic prediction. CD34+ cells grown in 5%, 14%, and 21% O_2_ also exhibited increased accumulation of intracellular reactive O_2_ species, a marker for oxidative stress (Figure 6E). Given the oxygen-dependent effects on reactive O_2_ species and mitochondrial gene expression (Figure 5D), we next sought to examine the O_2_ associated metabolic properties of HSCs/HPCs using the Seahorse metabolic flux analyzer. ATP rate assays showed that CD34+ cells generate significantly more ATP from glycolysis in lower O_2_ tensions (1%, 3%, and 5%) compared to 14% and 21%, at the expense of ATP generated from oxidative phosphorylation (Figure 6F). Cells grown in 1% had the lowest ratio of oxidative phosphorylation to glycolysis compared to all other expansion conditions (Figure 6F). To further examine the oxidative phosphorylation pathway, we examined the effects of inducing acute stress using the mitochondrial stress test (Figure S4A). CD34+ cells exhibited oxygen-dose dependent effects on mitochondrial respiration, with cells grown at 1% O_2_ exhibiting significantly lower basal and maximal respiration rates as well as lower spare respiratory capacity compared to 14% and 21% (Figure 6G-H, Figure S4A).

These data show that CD34+ cells grown at higher O_2_ tensions are more metabolically active, more proliferative, and accumulate more stress related molecular properties compared to those maintained in lower O_2_. Thus, we hypothesized that CD34+ cells in low physiologic O_2_ are more resistant to acute toxicities. To test this, we treated CD34+ cells with the broadly cytotoxic chemotherapy cytarabine in variable O_2_ levels. CD34+ cells grown in low O_2_ tensions, particularly 1% oxygen, were highly resistant to cytarabine induced cell death compared to higher O_2_ levels, as indicated by a significantly higher percentage of viable cell number after 2 days of treatment compared to vehicle (Figure 6I) and higher expansion over time in the presence of cytarabine (Figure S4B). These data provide insights into hematopoietic toxicities following chemotherapy and resistance of malignant hematopoietic cells to treatment.

### Hematopoietic mTOR is O_2_ dependent

Several genes that exhibit stabilized expression in low O_2_ tensions in HPC populations are linked to mTORc signaling (Figure 5D-E) which regulates processes critical to HSC/HPC fate including mitochondrial function, biogenesis, and degradation^42^. These include repressors of mTORc signaling (*BNIP3* and *DDIT4*)^37,38,43^ and genes downstream of mTORc (*LDHA* and *ENO1*)^40,41^. Interestingly, we found that the mTORc inhibitor Rapamycin partially reversed the O_2_ dependent expansion of total nucleated cells (Figure 6J) and led to maintenance of higher proportions of CD38-cells in high O_2_. Thus, inhibition of the O_2_-dependent mTORc partially rescued low O_2_ maintenance of the CD34+ cells enriched for potent HSCs/HPCs (Figure 6K). This is supported by work that demonstrated that short-term *ex vivo* Rapamycin treatment increases long-term and secondary repopulating capacities of human CD34+ cells expanded in ambient air^44^.

### O_2_ dependent transcripts predict engraftment outcomes

The O_2_-dependent, potency associated gene *PRSS2* (Figure 7A-B) encodes a Trypsin protein that has been primarily studied in the pancreas as a key digestive enzyme^35^. Its role in HSC/HPC biology is unclear, though its hematopoietic expression has been reported^36^. We examined *PRSS2* expression in HSC/HPC populations by sequencing immunophenotypically defined HSC/HPC subpopulations from freshly isolated CBUs and found that *PRSS2* is most highly expressed in MLPs followed by HSCs (Figure 7C, Table S3). In independently published data, HSCs exhibit significantly higher expression of *PRSS2* compared to mature HPC populations (Figure 7D). We hypothesized that *PRSS2* might be a transcriptional marker of HSC potency. To test this, we performed bulk RNA-seq on HSCs from biologically distinct CBUs with known engraftment outcomes in mouse models of transplantation (Figure 7E; Table S4,S5), modeling the CD34+ and HSC transcriptomes as a function of SCID repopulating cell (SRC) frequencies. This data revealed that high *PRSS2* expression in HSCs accurately predicts high SRC frequency (Figure 7F). Notably, the mTORc regulator *DDIT4* also trended toward higher expression in CBUs with higher SRC frequencies, but only in the broader CD34+ population (Figure S5). Together, these data showed that specific O_2_ dependent genes mark potent HSCs/HPCs in context of transplantation.

**Figure 7.**
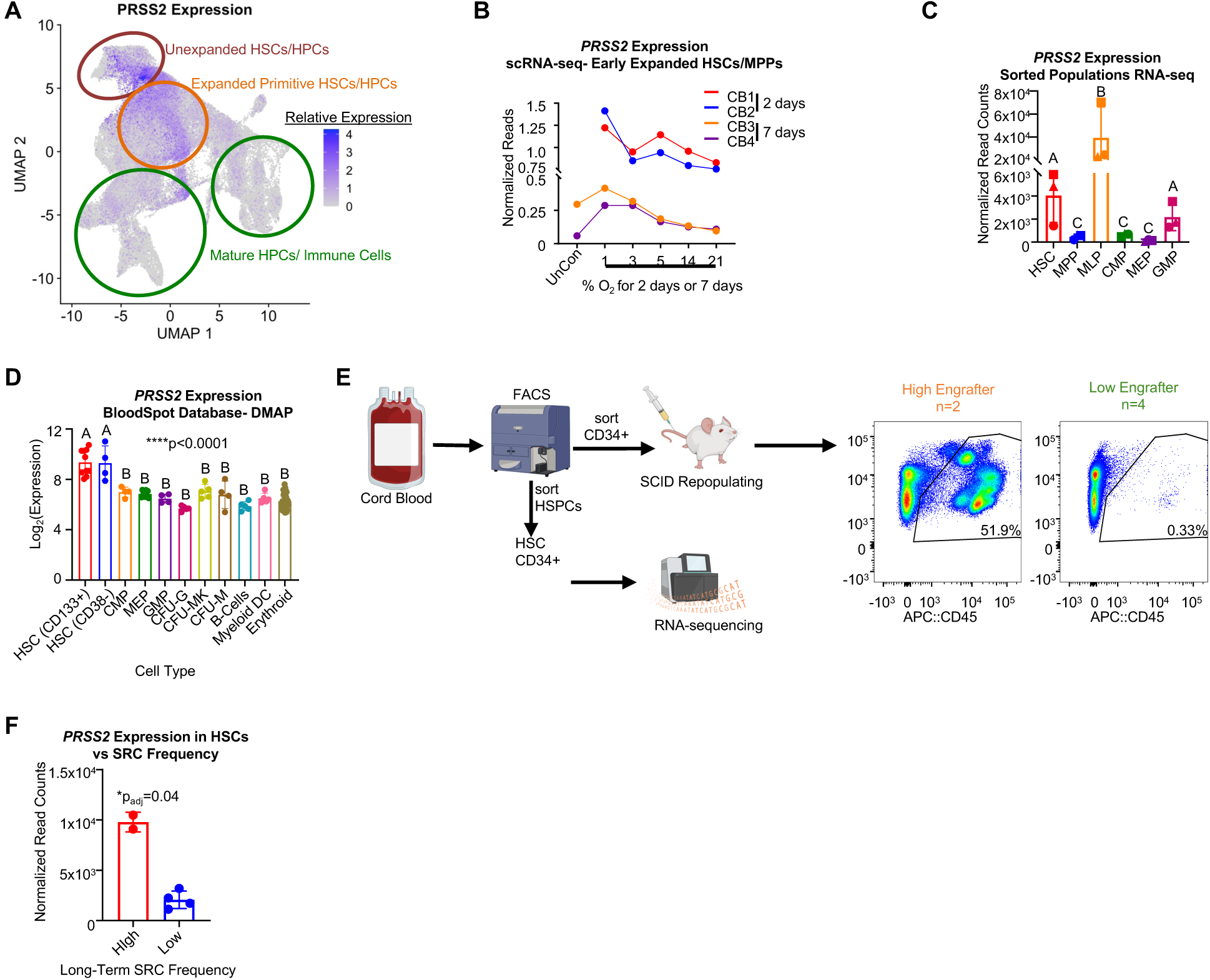
*PRSS2* predicts long-term engraftment of HSCs. A) Feature plot showing *PRSS2* expression in primitive HSCs/HPCs clusters from scRNA-seq B) Expression of *PRSS2* in different O_2_ tensions in HSCs/MPPs by scRNA-seq. C) Expression of *PRSS2* in immunophenotypically defined and sorted HSCs/HPCs by bulk RNA-seq. D) Expression of *PRSS2* in a HSC/HPC populations from a publicly available database. E) *PRSS2* expression in HSCs correlated with long-term engraftment capacity in a mouse model of human CB transplantation. DESeq2 used for significance of differential expression analysis. Stats: C-D) one-way ANOVA (indicated by p-value) with post hoc comparisons indicated by compact letter display (in compact letter display, groups that are not statistically different are indicated by matching letters). F) DESeq2 statistical analysis. UnCon = Unmanipulated control (input).

## Discussion

HSCs/HPCs are exposed to a wide range of O_2_ tensions during their physiologic lifespan and when being studied or utilized *ex vivo*^4–9^. Here we showed that HSCs/HPCs are acutely sensitive to physiologic O_2_ with regards to immunophenotype, self-renewal, and differentiation properties as well as *in vivo* potency and molecular profiles, particularly stress responses. All together, our data suggest that physiologic O_2_ is a critical extrinsic factor that affects the growth and function of HSCs/HPCs through regulation of stress associated molecular pathways. Multipotent hematopoietic cells remain the most quiescent, show the lowest biochemical signs of stress accumulation, and express genes associated with stem cell properties in low physiologic O_2_ tensions (1-3%), though their proliferative capacity is very specific to the specific cell subpopulation with HSC frequency highest in 5% and MPP frequency highest in 1-3%. When exposed to higher physiologic and extra physiologic tensions, HSCs expand poorly, suggesting that they are differentiating. Mature HPCs expand rapidly in higher O_2_ tensions and show properties associated with higher metabolism, rapid proliferation, and stress accumulation. Thus, HSCs/HPCs may utilize local BM O_2_ tensions to maintain the pool of HSCs or rapidly initiate differentiation and expansion of HPCs in response to different homeostatic or pathogenic states. The high O_2_ tensions these cells experience outside the BM likely intensifies HPC tendencies to proliferate, which would drive expansion of lineage defined progenitors in response to infection or wounding. Future studies should examine the O_2_ dependency of other cells of hematopoietic origin like mature lymphocytes.

Our findings have immediate translational implications. First, malignant hematopoietic cells are also exposed to variable O_2_ levels and have been shown to be affected by hypoxia. We infer from our data that O_2_-dependent gene programs are partially responsible for chemotherapeutic resistance and can use our transcriptomic profile to identify targets for treatment sensitization. Additionally, hematopoietic cell therapies including CB transplantation are a critical tool for treating patients with hematologic disorders and some non-hematologic disorders. A limiting factor in the efficacy of transplantation is the number of functionally potent HSCs/HPCs that can be delivered to the patient^18–20^. Here we have identified O_2_ dependent genes that mark potent HSCs/HPCs, including *PRSS2* and *DDIT4*, which could aid in the selection of the optimal donor units that are most likely to achieve favorable outcomes.

Further, oxygen-dependent gene programs represent therapeutic targets to improve the potency to regulate HSC/HPC responses to stress. Finally, understanding which conditions are best for use in *ex vivo* proliferation allows us to improve existing clinical protocols for hematopoietic cell therapies that require *ex vivo* manipulation, including CB expansion, gene editing, and the derivation of off-the-shelf immune effector therapies from HSCs. Previous studies have shown that human and mouse HSCs are best cultured in low O_2_ levels compared to higher O_2_ tensions^23–26^, but there are discrepancies about how low that tension should be. Our study provides insights into the nuanced biological O_2_ dependencies by examining source specific and cell type specific effects of different oxygenation. For example, our data suggests that very low physiologic O_2_ (1%) is ideal for maintenance of functionally potent CB HSCs and early HPCs, while mid-range O_2_ (3-5%) may be best for *ex vivo* expansion of early repopulating cells. Understanding the best conditions in which to expand HSCs/HPCs, as well as the best conditions to improve HPC commitment to target lineages will be critical for the improvement of these therapies. We demonstrated that BM and mPB are also sensitive to different O_2_ levels, though to different extents in different populations. Future work should optimize the leveraging of O_2_ dependencies in these donor sources for improved therapeutics.

In all, this study provides insight into how physiologic levels of O_2_ may drive changes in hematopoietic cell function during homeostasis and stress and how this can be leveraged to improve HSC/HPC potential for therapies.

## Acknowledgments

The authors thank Dr. Kelvin Lee for helpful feedback throughout the course of the study. This work was supported by Public Health Service Grants from the NIH: U54 DK106846 (MC), K99 HL166790 (JR), R00 HL166790 (JR), as well as an IU Simon Comprehensive Cancer Center Trainee Pilot Grant (JR) and an Indiana Clinical and Translational Sciences Institute Basic Research Grant (MC). We thank members of the Indiana University Melvin and Bren Simon Cancer Center Flow Cytometry Resource Facility, which is supported in part by P30 CA082709, U54 DK106846, and 1S10D012270. Sequencing was carried out in the Center for Medical Genomics at Indiana University School of Medicine, which is partially supported by the Indiana University Grand Challenges Precision Health Initiative. Mouse studies were supported by technical expertise from the *in vivo* Therapeutics Core at Indiana University School of Medicine, which is supported in part by the Indiana University Cooperative Center for Excellence in Hematology (U54 DK106846). We thank the Volunteer Donating Communities in Cleveland, OH, Atlanta, GA, and San Francisco, CA as the source of the Cord Blood Units used in this study.

## Author Contributions

Conceptualization, JR; Methodology, JR and MLC; Investigation JR, SG, LW, GW, AA, SC, WvH, MHK, and MLC; Formal Analysis JR and SG; Resources, JR, WvH, MHK, and MLC; Data Curation, JR; Writing-Original Draft, JR, MHK, and MLC; Writing-Review & Editing JR, SG, LW, GW, AA, SC, WvH, MHK, and MLC; Visualization, JR; Supervision, JR, MHK, and MLC; Funding Acquisition, JR, MHK, and MLC.

