## Supplemental Materials and Figures for "Local Oxygen Tension Dictates Hematopoietic Cell Growth and Potency"

**Detailed Methods:**

***CD34+ cells isolation or purchase:***

Fresh CB >= 60mL in volume and <55hrs since birth were collected in 35mL of Anticoagulant Citrate Phosphate Dextrose Solution and provided by or purchased from the Cleveland Cord Blood Center. Low density CB was collected using density centrifugation with Ficoll Paque (Cytiva). CD34+ cells were isolated using magnetic anti-CD34 monoclonal antibody and magnetic selection columns (Miltinyi). CD34+ cell pellets cryopreserved in 90% FBS + 10% DMSO freezing solution and stored in liquid nitrogen. Frozen CD34+ cells derived from three distinct human bone marrow donors were purchased from STEMCELL. Frozen CD34+ cells derived from three distinct mobilized peripheral blood donors were purchased from the Fred Hutch Cooperative Center for Excellence in Hematology Hematopoietic Cell Procurement and Processing Core. Prior to use in assays, CD34+ cells were thawed rapidly by warming in a 37°C water bath with gentle agitation and DMSO was washed out.

***Immunophenotyping by Cell Surface Staining and Fluorescence-Activated Cell Sorting:***

$1x10^5$-$1x10^6$ freshly thawed CD34+ enriched cells or cells from CD34+ expansion were incubated in PBS with fluorophore conjugated antibodies targeting CD34, CD38, CD45RA, CD10, CD49f, CD90, and CD135. Cells were washed 2x with cold PBS and were fixed with 1% paraformaldehyde (Pierce) and analyzed using FACS. Positive and negative gates were drawn based on unstained and single stained controls. Cell subpopulations were defined as follows: HSC: CD34+CD38-CD45RA-CD49f+CD90+; MPP: CD34+CD38-CD45RA-CD49f-CD90-; MLP: CD34+CD38-CD45RA+CD10+; CMP: CD34+CD38+CD10-CD45RA-CD135+; MEP: CD34+CD38+CD10-CD45RA-CD135-; GMP: CD34+CD38+CD10-CD45RA+CD135+. For cell cycle analysis, cells were permeabilized and stained with KI67 and DAPI. Cell populations were annotated as follows: G0: CD34+ DAPI low KI67-; G1: CD34+ DAPI low KI67 int; S: CD34+ DAPI int KI67 int; G2: CD34+ DAPI high KI67 int; M: CD34+ DAPI high KI67 high. For ROS analysis, cells were incubated with the cell permeant ROS detector CM-H2DCFDA (ThermoFisher) diluted to 5μM at 37°C for 30 minutes, washed twice, and analyzed immediately.

***Ex Vivo Expansion Assays:***

Thawed CD34+ cells were grown at a density of $5x10^4$ cells/mL in 1mL SFEM II (STEMCELL Technologies) supplemented with 100ng/mL recombinant human thrombopoietin (TPO), 100ng/mL recombinant human Stem cell factor (SCF), and 100ng/mL recombinant human FLT3 ligand (FLT3L) (R&D Technologies). Cultures were incubated at 37°C in a humidified incubator kept at 5% $CO_2$ and the $O_2$ level indicated by the experiment. These tensions were 1%, 3%, 5%, 14%, and 21% $O_2$. On day 4 after plating, fresh media containing the same growth factors that had been equilibrated in the appropriate oxygen tension overnight was added to a total volume of 2mL. On day 7 after plating, cells were collected, nucleated cellularity was counted by Trypan Blue staining, and expanded cells were used for immunophenotyping, CFU assays, or in vivo transplantation in mice.

***Ex Vivo Colony Forming Unit (CFU) Assays:***

CFU-GM, BFU-E, and CFU-GEMM colony numbers were derived by plating 250 freshly thawed CD34+ cells or 250 cells from CD34+ expansion assays in 1% methylcellulose/Iscove's Modified Dulbeccos Medium (IMDM) with 30% FBS (Corning), 1 U/mL recombinant human (rh) Epogen (EPO) (Amgen), 10ng/mL rh Interleukin-3 (IL-3) (R&D Systems), 50ng rh Stem cell factor (SCF) (R&D Systems), 10ng rh Granulocyte/macrophage colony stimulating factor (GM-CSF) (R&D Systems), 2mM L-glutamine, and 0.02mM 2-Mercaptoethanol. Cultures were incubated at 37°C in a humidified environment containing 5% $CO_2$ and the $O_2$ level indicated by the experiment. For direct analysis of oxygen tension effects on CFU, this was 1%, 3%, 5%, 14%, and 21% $O_2$. For post expansion CFU numbers, this was 5% $O_2$. Colonies were scored by manual counting 12 days after plating using an inverted microscope with phase contrast (Nikon) with a 4x objective.

***In Vivo Model of Oxygen Dependent HSC/HPC Potency:***

7 days prior to transplantation, CD34+ cells from 3 distinct CBUs were expanded as described above in 1%, 3%, 5%, 14%, or 21% $O_2$. 24 hours prior to transplantation, NSG mice were given a sublethal (350gy) whole body dose of gamma radiation. Irradiated mice were transplanted by tail vein injection with 2.5x104 freshly thawed unmanipulated CD34+ enriched cells or 2.5x104 CD34+ cells that had been plated in expansion plus all expanded progeny (200uL per injection). Each group (CBU + oxygen tension) contained 3 mice, for a total of n=9 mice and n=3 CBUs per expansion condition. 2 weeks, 6 weeks, 10 weeks, and 14 weeks following transplantation, <= 50uL of peripheral blood was collected from each mouse in EDTA tubes or tubes containing heparin by submandibular bleeding. 16 weeks following transplantation, mice were euthanized and bone marrow was harvested by flushing. Equivalent cell numbers from each primary recipient mouse bone marrow were pooled per group (CBU + oxygen tension) and 1x106 pooled bone marrow cells were transplanted to secondary recipient mice that had received a

350gy dose of radiation 24 hours prior. A total of n=5 mice and n=3 CBUs were used for secondary transplantations. Secondary mice were euthanized 16 weeks after transplantation and bone marrow was harvested by flushing.

***Analysis of Peripheral Blood and Bone Marrow from In Vivo Studies:***
1x106 flushed BM or red blood cell depleted PB cells were stained with fluorophore conjugated antibodies: at week 2 anti-CD45, anti-CD66b, and anti CD15 for neutrophil detection; at all other time points anti-CD45, anti-CD33, anti-CD19, and anti-CD3 for human chimerism and myeloid/lymphoid ratio analysis. Cells were washed 3 times, fixed, and analyzed by FACS.

***Dissolved Oxygen Measurements of CB:***
Dissolved oxygen was measured from plasma of pelleted cord blood using an Orion Star A323 Dissolve Oxygen Portable Meter. A total of n=70 distinct CBUs were measured.

***Single Cell RNA-sequencing Library Preparation by PIP-seq:***
CD34+ enriched cells were harvested following thawing, or were expanded as described above and were collected after 2 days or 7 days of expansion (2 CBUs were expanded for 2 days, 2 CBUs were harvested as input and were expanded for 7 days). Cells harvested after 2 days had media refreshed with appropriate $O_2$ equilibrated media one day following plating and those expanded for 7 days had media refreshed every 2 days, to maintain fresh growth factors. On the day of harvest, cells expanded in low oxygen (1%, 3%, and 5%) were harvested in a hypoxia chamber equilibrated to 3% oxygen, while unmanipulated cells and those expanded at high oxygen tensions (14% and 21%) were harvested in an ambient air biosafety cabinet. Cells were counted and equivalent numbers of cells were incubated with cell hashing antibodies (BioLegend). A different cell hashing antibody was used for each oxygen tension and one additional was used for the unmanipulated samples. Cells were washed. Equivalent numbers of cells from each condition were pooled into one tube for each CBU. At this point cells were processed in ambient air but were kept on ice as much as possible. Total exposure to ambient air prior to cell lysis was ~90 minutes. Using the PIPseq T20 kit (Fluent Biosciences), $5x10^4$ cells from each CBU was subjected to single cell barcoding using particle emulsions. Cells were then lysed and single cell gene expression libraries and cell hashtag libraries were prepared according to manufacturer’s recommendations. Cell hashing libraries were spiked in at a 1:10 ratio to the gene expression libraries and samples were submitted for sequencing to the Indiana University Center for Medical Genomics. 2-day expanded samples were sequenced for

2.5billion paired end reads on a NovaSeq 6000. Input and 7-day expanded samples were sequenced for 2.5billion paired end reads on a NovaSeqX Plus. Raw fastq files passing quality filters were obtained from the Center for further analysis.

***scRNA-sequencing Analysis:***

Raw fastq files were aligned to the human genome and assigned to cell barcodes and cell hashtags using PIPSeeker software (FluentBiosciences). Data was imported into R using Seurat. Remaining analyses were performed using Seurat v5. Data were normalized using SCTransform and apoptotic cells were controlled for using mitochondrial read percentages. After observing clear batch effects, the four different CBUs were integrated using Harmony. Cells lacking a hashtag or containing multiple hashtags were removed from analysis. Principal components were determined computationally. Clustering resolution was determined by comparing several different resolutions and selecting 1.0, which best separated several clusters of interest.  Annotation was performed by finding markers unique to each cluster compared to all other clusters pooled, as well as by direct comparison of each cluster to every other cluster, and then by manual annotation. Full details on genes used to make annotation decisions are found in the publicly available R Script on Mendeley. Markers that define each annotated cluster are found in Supplemental Table 6.

***Seahorse Metabolic Analysis:***

CD34+ enriched cells were expanded as described above. Cells were washed twice with de-OCR media (Agilent Seahorse XFe RPMI media supplemented with 10mM Glucose, 1mM Pyruvate, and 2mM Glutamine). 5x104 cells were adhered to the plate using CellTak (Corning). Plates were subjected to either the ATP Rate Assay or the Mitochondrial Stress Test Assay on a Seahorse XFe96 metabolic flux analyzer, according to manufacturer recommendations. FCCP was used at 1μM.

***Chemosensitivity Assays with Cytarabine:***

CD34+ enriched cells from CB were expanded as described above, in the presence or absence (DMSO vehicle) of 0.02μM Cytarabine. Cells were counted with Trypan Blue,

***Western Blot Protein Expression Analysis:***

$1x10^5$ cells were lysed for 30 minutes on ice using RIPA buffer (ThermoFisher) containing Roche EDTA-free protease inhibitor tablets and cleared by high speed centrifugation. Lysates were run on a 4-20% denaturing Mini-Protean TGX Gel (BioRad). Proteins were transferred to

0.45µm nitrocellulose membranes. Membranes were blocked in Blocking Buffer + 0.05% Tween-20, and blotted with anti-ENO1, anti-BNIP3, anti-LDHA, or anti-ACTB followed by anti-rabbit-HRP secondary. Bands were detected with Femtoluminescent substrate (Pierce) and imaged on a ChemiDocX (BioRad).

***Rapamycin Treatment of CB:***
CD34+ enriched cells from CB were expanded as described above, in the presence or absence of 25nM Rapamycin (DMSO vehicle control). Cells were analyzed for immunophenotype.

***Immunophenotypic HSCs/HPCs transcriptomics:***
CD34+ enriched cells were thawed as described above and stained for immunophenotyping as described above. HSCs, MPPs, MLPs, CMPs, MEPs, and GMPs were sorted by FACS from three distinct CBUs and collected directly in RLT Lysis Buffer. After collection of >= $5x10^2$ cells, tubes were vortexed for 30 seconds, snap frozen, and stored in -80°C.

***Long Term Engraftment/ SCID Repopulating Cell Frequency transcriptomic model:***
Immunophenotypically defined HSCs or CD34+ cells were stained with fluorescently conjugated antibodies as described above and were sorted by FACS from three distinct CBUs and collected directly in RLT Lysis Buffer. After collection of >= $5x10^2$ cells, tubes were vortexed for 30 seconds, snap frozen, and stored in -80°C. The remainder of CD34+ enriched cells from the same CBUs were used for limiting dilution SCID repopulating assay. 24 hours prior to transplantation, NSG mice were given a sublethal (350rads) whole body dose of radiation. Irradiated mice were transplanted by tail vein injection with varying cell doses of $1x10^3$, $5x10^3$, or $10x10^3$ CD34+ enriched cells, 200uL per injection. Each group (cord blood unit + cell dose) contained 7-8 mice.

20 weeks following transplantation, mice were euthanized and BM cells were harvested by flushing the right femur. BM cells were counted. $3x10^6$ cells were then stained with APC anti-humanCD34 to measure human chimerism. For SCID repopulating cell (SRC) frequency calculations, mice with > 0.2% human chimerism in the bone marrow of recipient mice at week 20 after transplantation were considered responders. This value is 2.15-fold higher than background observed in an untransplanted control. SRC frequencies were determined using Extreme Limiting Dilution Analysis (ELDA) software available from Walter+Eliza Hall Bioinformatics (https://bioinf.wehi.edu.au/software/elda/) (1). SRC frequencies were back-

calculated based on CD34+ purity of the CD34+ enriched cells used for transplantation to account for differences.

***RNA Extraction and Library Prep for Bulk Transcriptomics models:***

For collection of RNA, frozen tubes were gently thawed at 4°C. Tubes were then vortexed again for 30 seconds and allowed to sit at room temperature for 5 min to ensure complete lysis. RNA was then harvested per manufacturer instructions using the Qiagen RNeasy Micro Plus Kit. High quality RNA (RIN => 8) was used to prepare amplified libraries either with the Indiana University Center for Medical Genomics using the SMART-Seq v4 Ultra Low Input RNA Kit (Takara) or in lab with the NEBNext Ultra Low Input RNA Sequencing Kit (NEB). mRNA libraries were submitted to the Indiana University Center for Medical Genomics and were sequenced on a NovaSeq v1.5 S4 (200 cycles kit) for paired end sequencing. Raw fastq files passing quality filters were obtained from the Center for further analysis.

***Bulk RNA-sequencing Analysis***

Fastq files were analyzed for quality using FastQC. Reads were trimmed of adapters and low-quality reads using Cutadapt with the arguments -a CTGTCTCTTATA -A CTGTCTCTTATA --nextseq-trim=20 --minimum-length 50. Raw genome files downloaded from GENCODE (GRch38 version 41) were used as input to generate genome indices with STAR alignment software using the arguments –sjdbOverhang 99. Trimmed reads were aligned to the genome using STAR with the arguments --readFilesCommand zcat --outSAMtype BAM SortedByCoordinate --outSAMunmapped Within. Aligned reads were assigned to exons and annotated by gene using HTSeq with the arguments -s no -r pos -f bam. Alternatively to STAR and HTSeq, genes were aligned to the human genome and assigned to genes using Kallisto. Gene counts were normalized and differential expression was analyzed using DESeq2 R package. Sex of the baby that the CBU came from was inferred by expression of *XIST* (female specific transcription) and *DDX3Y* (male specific transcription) so that sex can be controlled for in the model where appropriate. The design used to examine differential expression between immunophenotypically defined HSCs/HPCs was ~CBU + celltype. The design used to examine differential expression between early engrafted/ homed cells and input was ~CBU + sampleType. The design used to examine differential expression between High engraftment/SRCs and Low engraftment/SRCs was ~Sex + Response. Gene set enrichment analysis was performed using test statistics generated by DESeq2 against curated datasets from MSigDB c2, c5, and c6 (downloaded from Walter+Eliza Hall Bioinformatics resource https://bioinf.wehi.edu.au/MSigDB/v6.1/) with fgsea R package with the arguments minSize = 50, maxSize = 500 and adjusted p-value cutoff of 0.05.

### *Statistical Analyses for Non-Sequencing Approaches*

All appropriate statistical analyses were determined prior to performing experiments. For nearly all *ex vivo* studies one-way ANOVA was performed with post hoc Tukey or Sidak multiple comparisons. For Rapamycin treatment studies, two-way Anova was performed with post hoc Tukey's multiple comparisons for Vehicle vs Treated. For in vivo studies, mixed-effects linear modeling was performed to assess differences in human chimerism corrected for CBU, using CBU as a randomly sampled variable and oxygen tension as the fixed variable. Pairwise comparisons used Tukey corrections.

**<u>Description of Supplemental Tables:</u>**

**Supplemental Table 1. scRNA-seq summary.** Separate excel file that contains the most pertinent data from the scRNA-seq data that was discussed in this study. scRNA-seq raw data and processed Seurat object will also be made publicly available.

**Supplemental Table 2. Distribution of cells in each annotated cluster as a fraction of total sequenced cells in each sample.** Separate excel file showing the frequencies of cells found in the indicated annotated cell cluster from each sample (i.e., Input, 1% $O_2$ 2 days, 1% $O_2$ 7 days, 3% $O_2$ 2 days, etc.) calculated as a proportion of the total cells in that sample. Red-Blue coloring is indicative of column z-scores, where deep red indicates a high proportion of cells found in that cluster and dark blue indicates a low proportion of cells found in that cluster.

**Supplemental Table 3. RNA-seq summary for sorted HSPCs.** Separate excel file that contains a sheet for normalized read counts for all samples and for all differential expression analyses discussed in this study.

**Supplemental Table 4. SRC calculations for transcriptomic analysis of engraftable HSCs.** Separate excel file showing the values input into the ELDA model (https://bioinf.wehi.edu.au/software/elda/index.html)(1) and the calculated lower and upper limits for SCID repopulating cell (SRC) frequencies as well as the estimated SRC frequency for each CBU. Mean value of estimated SRC frequencies for all samples was 8.E-4. High SRC frequency was assumed for all CBUs that had greater SRC frequency than the mean.

**Supplemental Table 5. RNA-seq summary for SRC model.** Separate excel file that contains a sheet for normalized read counts for all samples and for differential expression analyses results.

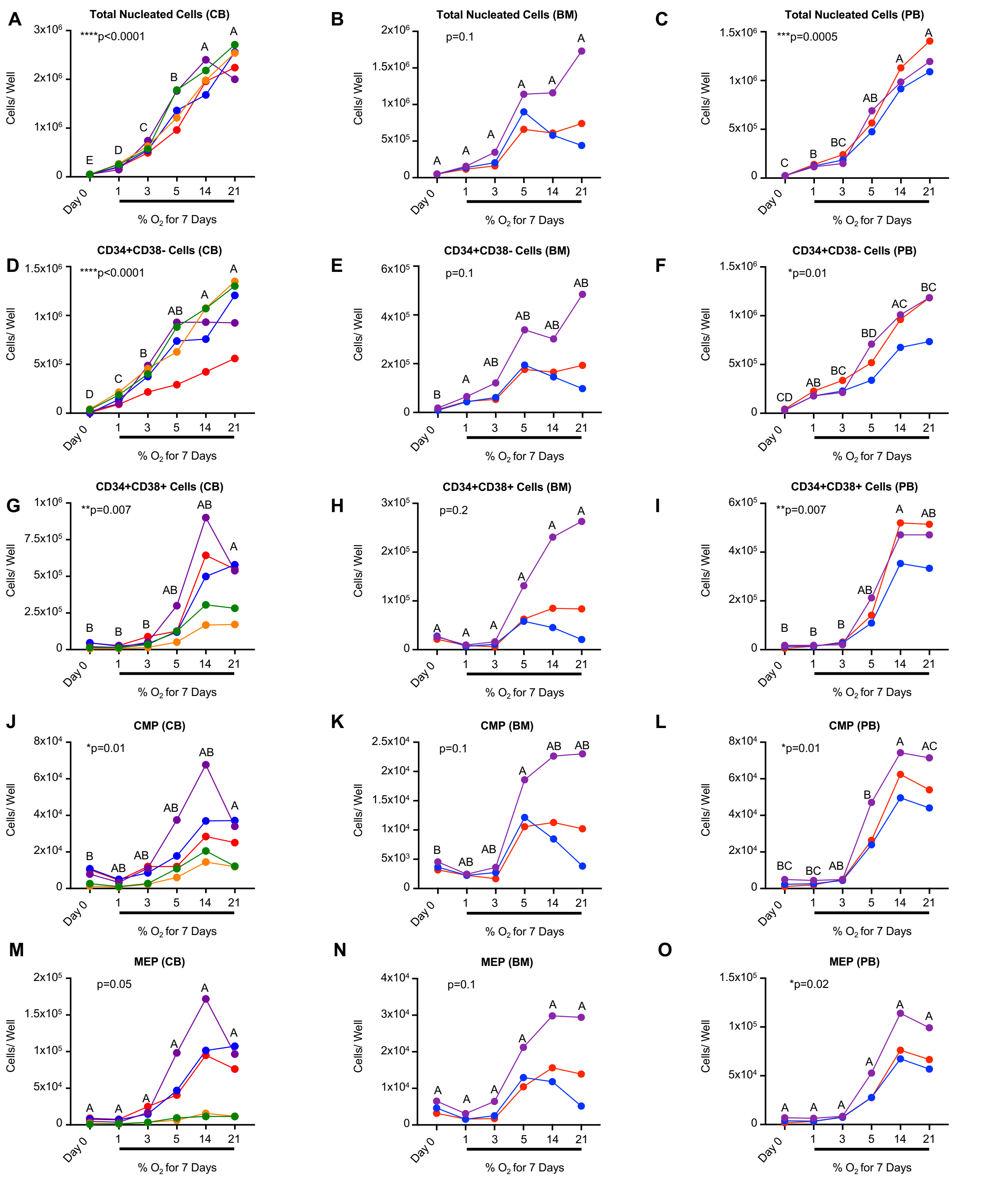


**Supplemental Figure 1 (Related to Figure 2-3).** A-O) Cord blood (CB), bone marrow (BM), or mobilized peripheral blood (PB) CD34+ cells were expanded in serum free media with growth factors for 7 days in the indicated oxygen tensions. Cells were then analyzed for enumeration of the indicated immunophenotypically defined population by flow cytometry (CB n=5, BM/PB n=3). Stats: one-way ANOVA (indicated by p-value) with post hoc comparisons indicated by compact letter display (in compact letter display, groups that are not statistically different are indicated by matching letters). Each point/color indicates distinct biological donor sources as in Figures 2/3.

**A**

| | input_7d | 1perc_2d | 1perc_7d | 3perc_2d | 3perc_7d | 5perc_2d | 5perc_7d | 14perc_2d | 14perc_7d | 21perc_2d | 21perc_7d |
|---|---|---|---|---|---|---|---|---|---|---|---|
| 0 | 132 | 1290 | 375 | 1307 | 356 | 1213 | 376 | 1245 | 259 | 1212 | 239 |
| 1 | 29 | 268 | 237 | 1398 | 234 | 1273 | 317 | 1362 | 252 | 1462 | 244 |
| 2 | 178 | 471 | 508 | 617 | 561 | 591 | 576 | 618 | 538 | 538 | 523 |
| 3 | 50 | 692 | 322 | 665 | 367 | 676 | 333 | 851 | 348 | 809 | 363 |
| 4 | 29 | 503 | 259 | 917 | 270 | 714 | 325 | 677 | 366 | 828 | 389 |
| 5 | 1822 | 2784 | 24 | 221 | 11 | 163 | 9 | 271 | 7 | 187 | 5 |
| 6 | 24 | 1868 | 171 | 656 | 157 | 564 | 159 | 758 | 160 | 672 | 159 |
| 7 | 445 | 350 | 225 | 571 | 273 | 574 | 362 | 529 | 494 | 549 | 495 |
| 8 | 7 | 560 | 122 | 790 | 118 | 675 | 108 | 769 | 105 | 763 | 121 |
| 9 | 25 | 273 | 72 | 770 | 89 | 730 | 95 | 883 | 72 | 789 | 91 |
| 10 | 82 | 175 | 278 | 232 | 321 | 191 | 393 | 224 | 535 | 220 | 609 |
| 11 | 35 | 55 | 213 | 172 | 288 | 125 | 519 | 161 | 640 | 162 | 672 |
| 12 | 759 | 22 | 215 | 128 | 225 | 97 | 202 | 76 | 316 | 127 | 330 |
| 13 | 20 | 212 | 58 | 517 | 70 | 563 | 93 | 483 | 95 | 574 | 101 |
| 14 | 2431 | 98 | 0 | 6 | 0 | 12 | 0 | 11 | 0 | 12 | 0 |
| 15 | 41 | 364 | 84 | 368 | 86 | 295 | 85 | 332 | 69 | 298 | 59 |
| 16 | 28 | 141 | 105 | 285 | 89 | 273 | 136 | 362 | 110 | 334 | 127 |
| 17 | 10 | 319 | 81 | 337 | 83 | 270 | 109 | 280 | 114 | 304 | 118 |
| 18 | 160 | 42 | 334 | 129 | 253 | 106 | 191 | 125 | 179 | 152 | 221 |
| 19 | 29 | 18 | 363 | 20 | 358 | 11 | 283 | 12 | 291 | 14 | 364 |
| 20 | 358 | 11 | 247 | 48 | 211 | 47 | 121 | 37 | 119 | 47 | 136 |
| 21 | 10 | 2 | 161 | 3 | 201 | 4 | 208 | 7 | 351 | 7 | 426 |
| 22 | 58 | 215 | 36 | 100 | 43 | 159 | 56 | 186 | 56 | 179 | 62 |
| 23 | 273 | 8 | 121 | 48 | 124 | 46 | 99 | 58 | 98 | 49 | 103 |
| 24 | 15 | 93 | 42 | 162 | 57 | 161 | 57 | 150 | 66 | 139 | 72 |
| 25 | 26 | 76 | 50 | 63 | 67 | 61 | 100 | 75 | 91 | 73 | 88 |
| 26 | 19 | 77 | 20 | 88 | 24 | 78 | 28 | 82 | 35 | 74 | 34 |
| Total | 7095 | 10987 | 4723 | 10618 | 4936 | 9672 | 5340 | 10624 | 5766 | 10574 | 6151 |

**B**

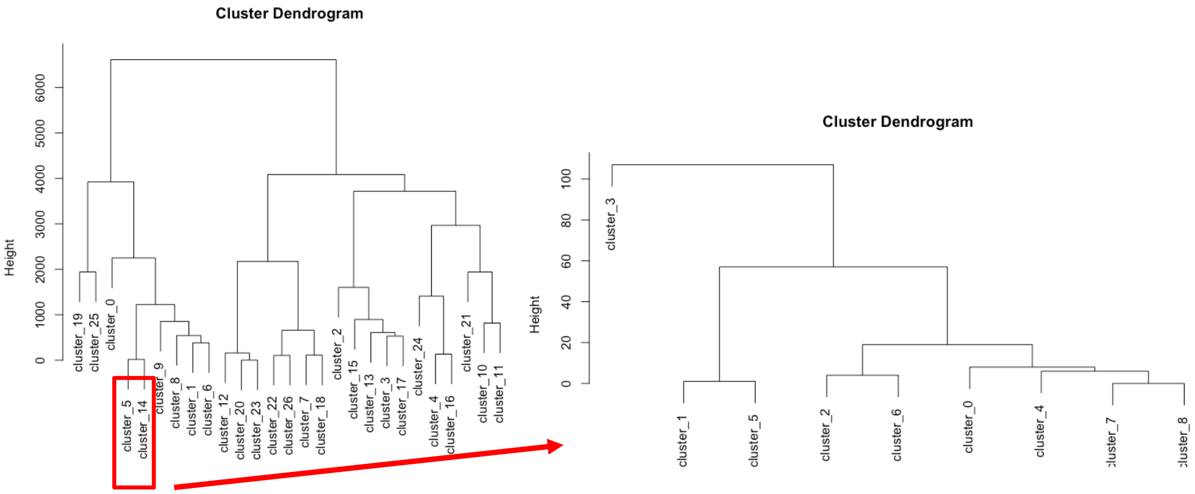


**Supplemental Figure 2 (Related to Figure 5).** A-B) CB CD34+ cells expanded in serum free media with growth factors for 2 days (n=2) or 7 days (n=2) in the indicated oxygen tensions, or unmanipulated input cells (n=2) were subjected to scRNA-seq using PIP-seq. A) Shown are the frequencies of cells found in the indicated initial cell cluster from each sample (i.e., Input, 1% $O_2$ 2 days, 1% $O_2$ 7 days, 3% $O_2$ 2 days, etc.) calculated as a proportion of the total cells in that sample. Red-Blue coloring is indicative of column z-scores, where deep red indicates a high proportion of cells found in that cluster and dark blue indicates a low proportion of cells found in that cluster. B) Dendrogram showing similarity scores between the different unannotated clusters. This dendrogram was used for direct comparison between clusters to perform manual annotation. Clusters 5 and 14 were subclustered due to most input cells being contained in these clusters to get a higher resolution of that sample.

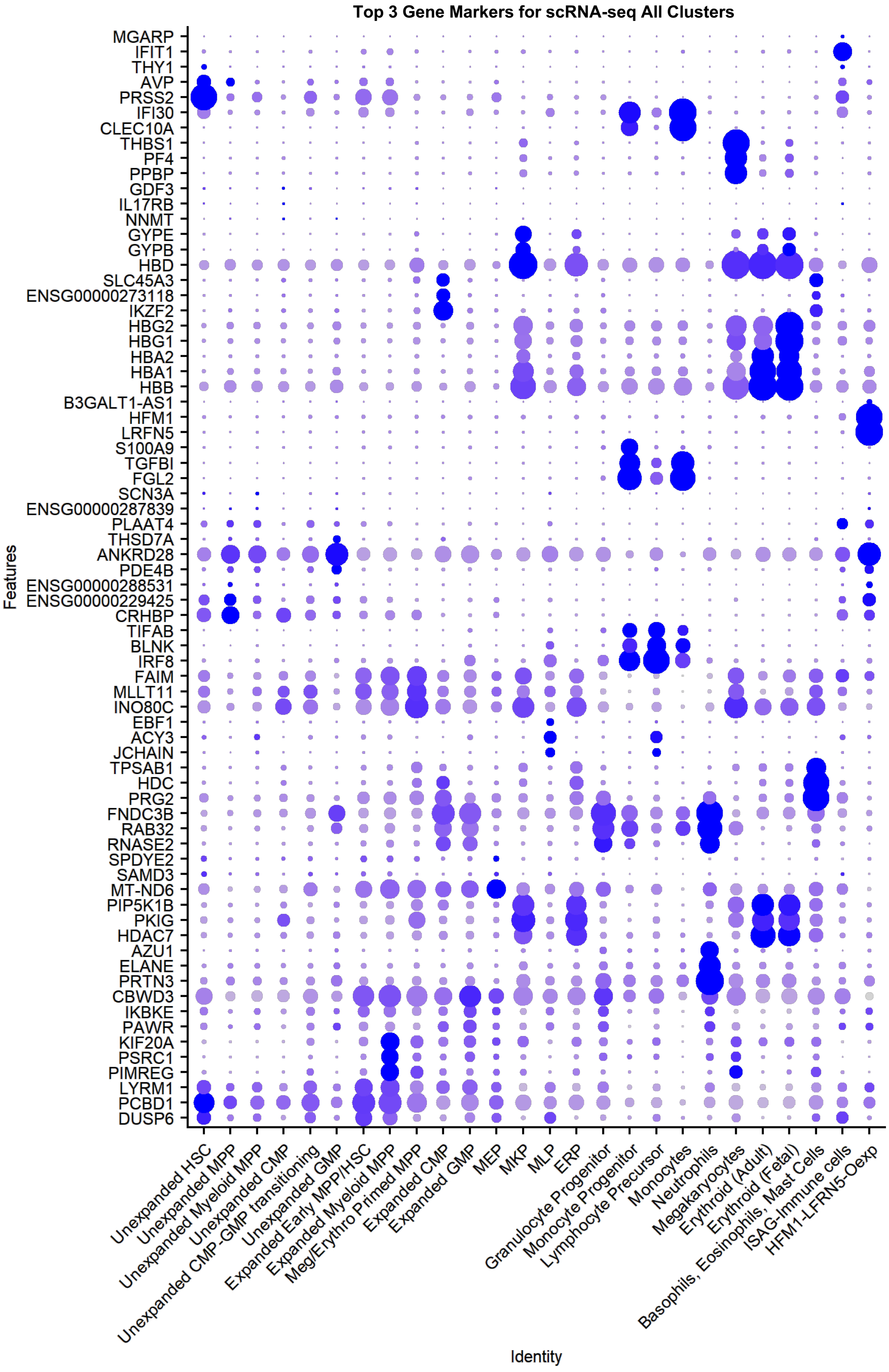


**Supplemental Figure 3 (Related to Figure 5).** Dot Plot showing top 3 gene markers used to annotate all cell clusters.

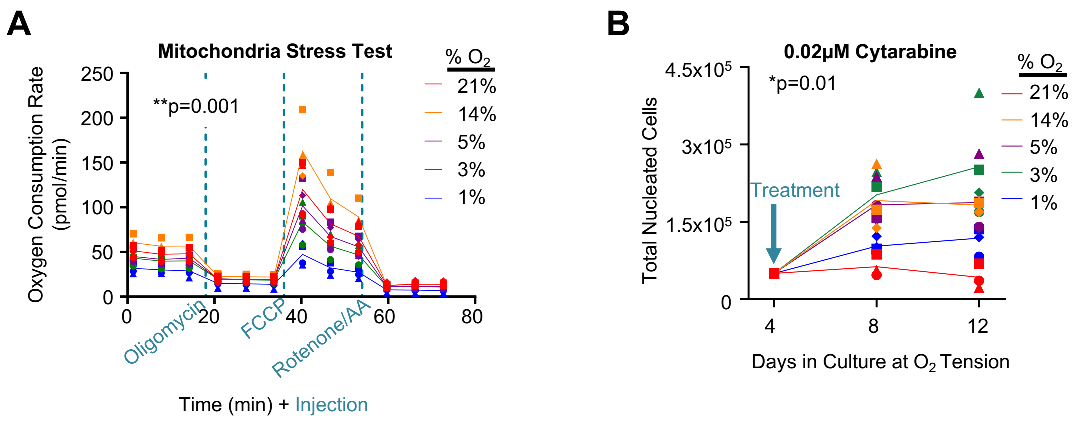


**Supplemental Figure 4 (Related to Figure 6).** A) Seahorse metabolic flux analysis after 7 days expansion using mitochondrial stress test (n=4) to show oxygen consumption rate following injections of Oligomycin, FCCP, and Rotenone/ Antimycin A. B) CD34+ cells were treated with 0.02μM cytarabine (n=3) for 8 days after equilibrating for 4 days in variable $O_2$ tensions and were counted for total nucleated cellularity. Stats: one-way ANOVA (indicated by p-value). Individual points indicate different CB donor units.

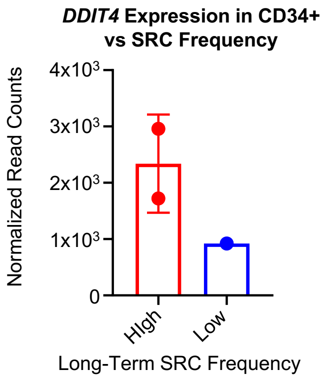


**Supplemental Figure 5 (Related to Figure 7).** DDIT expression in CD34+ cells correlated with long-term engraftment capacity in a mouse model of human cord blood transplantation. DESeq2 used for significance of differential expression analysis, not statistically significant due to low sample size.

**Supplemental Material References**

1. Y. Hu, G.K. Smyth, ELDA: extreme limiting dilution analysis for comparing depleted and enriched populations in stem cell and other assays. *J. Immunol. Methods* **347**, 70–78 (2009).
